# A Phytophthora nucleolar effector, Pi23226, targets to host ribosome biogenesis for necrotrophic cell death

**DOI:** 10.1101/2022.05.02.490323

**Authors:** Soeui Lee, Jaehwan Kim, Myung-Shin Kim, Cheol Woo Min, Sun Tae Kim, Sang-Bong Choi, Joo Hyun Lee, Doil Choi

**Affiliations:** Plant Immunity Research Center, Seoul National University, Seoul, 08826, Republic of Korea; Plant Genomics and Breeding Institute, Department of Agriculture, Forestry and Bioresources, College of Agriculture and Life Science, Seoul National University, Seoul, 08826, Republic of Korea; Interdisciplinary Programs in Agricultural Genomics, College of Agriculture and Life Science, Seoul National University, Seoul, 08826, Republic of Korea; Department of Plant Bioscience, Life and Industry Convergence Research Institute, Pusan National University, Miryang 50463, Republic of Korea; Division of Bioscience and Bioinformatics, Myongji University, Yongin 449–728, Republic of Korea

**Keywords:** *Phytophthora infestans*, hemibiotroph, RXLR effector, cell death, ribosome biogenesis, nucleolus, translation inhibition, plant immunity

## Abstract

Pathogen effectors target diverse subcellular organelles to manipulate the plant immune system. Although nucleolus has been emerged as a stress marker, and several effectors are localized in the nucleolus, the roles of nucleolar-targeted effectors remain elusive. In this study, we showed *Phytophthora infestans* infection of *Nicotiana benthamiana* results in nucleolar inflation during the transition from biotrophic to necrotrophic phase. Multiple *P. infestans* effectors were localized in the nucleolus: Pi23226 induced cell death in *Nicotiana benthamiana* and nucleolar inflation similar to that observed in the necrotrophic stage of infection, whereas its homolog Pi23015 and a deletion mutant (Pi23226ΔC) did not induce cell death or affect nucleolar size. RNA immunoprecipitation and iCLIP-seq analysis indicated that Pi23226 bound to the 3′-end of 25S rRNA precursors, resulting in the accumulation of unprocessed 27S pre-rRNAs. The nucleolar stress marker NAC082 was strongly upregulated under Pi23226-expressing conditions. Pi23226 subsequently inhibited global protein translation in host cells by interacting with ribosomes. Pi23226 enhanced *P. infestans* pathogenicity, indicating that Pi23226-induced ribosome malfunction and cell death was beneficial for pathogenesis in the host. Our results provide evidence for the molecular mechanism underlying RNA-binding effector activity in host ribosome biogenesis, and lead to new insights into the nucleolar action of effectors in pathogenesis.

## Introduction

Plants developed a unique innate immune system against pathogen infection because they lack an adaptive immune system (Jones and Dangl, 2006; Jones et al., 2016). Plant contains plasma membrane-localized pattern recognition receptors (PRRs) that recognize pathogen-associated molecular patterns (PAMPs) and initiate pattern-triggered immunity (PTI) (Chisholm et al., 2006). During pathogen invasion of host cells, specialized secreted molecules called effectors are delivered to modulate the host immune response and enhance pathogenicity. As a result of the long arms-race history between hosts and pathogens, plants have evolved polymorphic pathogen resistance (R) genes that contain a nucleotide-binding site and a leucine-rich repeat domain (NLR) to recognize cognate pathogen effectors. Effector recognition induces a more durable and robust defense response called effector-triggered immunity (ETI) (Cui et al., 2015). ETI often results in a hypersensitive cell death response (HR) that restricts further pathogen invasion at the infection site (Dangl et al., 2013).

Recent research has focused on completing the whole genome sequences of pathogens and understanding the molecular mechanisms of effector action in pathogenesis and host defense. RXLR-type effectors are ubiquitous in oomycete pathogens and are a major target for understanding their molecular actions in host cells (Birch et al., 2006; Dou and Zhou, 2012; He et al., 2020; Rehmany et al., 2005). After pathogen effectors are translocated into plant cells, they target diverse subcellular organelles to manipulate the plant immune system. For example, the *P. infestans* effector, Pi04314 modulates the activity of protein phosphatase I in the nucleus and suppresses JA- and SA-induced transcription of defense-related genes (Boevink et al., 2016). The *P. sojae* RXLR effector, PsAvh262 associates with endoplasmic reticulum (ER)-luminal binding immunoglobulin proteins and suppresses ER stress-related cell death (Jing et al., 2016).

The nucleolus is a eukaryotic subcellular organelle that has critical functions in ribosome biogenesis, initiating rDNA transcription, and processing pre-rRNAs to assemble mature rRNA-ribosomal protein translation complexes (Boisvert et al., 2007; Kalinina et al., 2018; Sáez-Vásquez and Delseny, 2019). RNA polymerase I mediates the transcription of rRNA genes, and nascent pre-rRNAs are processed by numerous ribonucleoproteins to produce 18S, 5.8S, and 25S rRNAs. The assembly of ribosome subunits requires more than 200 accessory factors known as ribosome biogenesis factors. Finally, mature rRNAs are assembled with ribosomal proteins to form small (40S) and large (60S) subunits in the nucleolus (Weis et al., 2015). Disruption of ribosome biogenesis factors causes severe defects in development. For example, homozygous knockout mutants of 40S ribosome biogenesis factors were embryo lethal in Arabidopsis (Harscoët et al., 2010; Missbach et al., 2013).

The nucleolus also has an emerging role in plant immunity against diverse pathogens. For example, a number of plant viral proteins target nucleolar methyltransferase fibrillarin (FIB) for the long-distance movement of viruses (Chang et al., 2016; Kim et al., 2007; Rajamäki and Valkonen, 2009). Plant nucleoli also respond to the invasion of bacterial, fungal, and oomycete pathogens (Seo et al., 2019). Under normal conditions, FIB2 interacts with Mediator subunit 19a (MED19a), a component of the Mediator complex, and functions as a negative regulator of immune-related gene expression. However, infection with the bacterial pathogen *Pseudomonas syringae* or treatment with Elf18 causes FIB2 dissociation from the complex and release the gene promoter to enhance the expression of immune-responsive genes. The *Hyaloperonospora arabidopsidis* RXLR effector HaRxL44 interacts with MED19a to modulate the interactions between transcriptional regulators and RNA polymerase II, which shifts the balance of transcription from SA-responsive to JA/ethylene-responsive genes, and enhances plant susceptibility (Caillaud et al., 2013). The *P. infestans* RXLR effector Pi04314 localizes in the host nucleus/nucleolus, which appears to be crucial for increasing pathogen colonization. Pi04314 interacts with phosphatase 1 catalytic (PP1c) isoforms, which are ubiquitous phosphatases in many cellular processes, and mimics PP1-interacting proteins to dephosphorylate key substrates in plant defense (Boevink et al., 2016). Although recent research has achieved significant progress in mapping the interactions of nuclear/nucleolar-localizing pathogen effectors, the molecular roles of nucleolar targeting effectors in pathogenicity and host defense are still elusive.

In this study, we observed the morphological changes of plant nucleoli during transition of the pathogen life cycle from biotrophic to necrotrophic stage. We also investigated the molecular activity of the *P. infestans* effector Pi23226, which is localized in the host nucleolus, in host ribosome biogenesis. We found that Pi23226 inflated the size of the nucleolus, subverted 25S pre-rRNA processing by binding immature 25S rRNA, and subsequently suppressed protein translation leading to host cell death. To our knowledge, Pi23226 is the first reported plant pathogen effector that directly targets and binds RNA. Our results provide new insights into the mechanisms of action of a nucleolar-localized pathogen effector that induces necrotic host cell death and enhances host susceptibility by subverting ribosome biogenesis and protein synthesis in the host cell.

## Results

### *P. infestans* infection leads to inflation of host nucleoli before the initiation of necrosis

*P. infestans* is a hemibiotrophic pathogen, and a necrosis occurs in colonized cells during a later stage of infection. We tracked the disease progress by inoculating *P. infestans* zoospores onto detached *N. benthamiana* leaves (500 spores/drop), and then collecting inoculated leaf tissues for 5 days after inoculation. Each collected sample was analyzed to determine the transcript levels of a biotrophic (IpIO) and necrotrophic (NPP1.1) marker (Kanneganti et al., 2007; Van West et al., 1998). IpIO expression dramatically increased at 2- and 3-days post-infection (dpi) compared to 0 dpi, and then subsequently decreased. By contrast, NPP1.1 expression sharply increased at 5 dpi (Figure 1B). These results suggested that the transition from biotrophic to necrotrophic phase occurred around 4 dpi in *N. benthamiana*.

**Figure 1.**
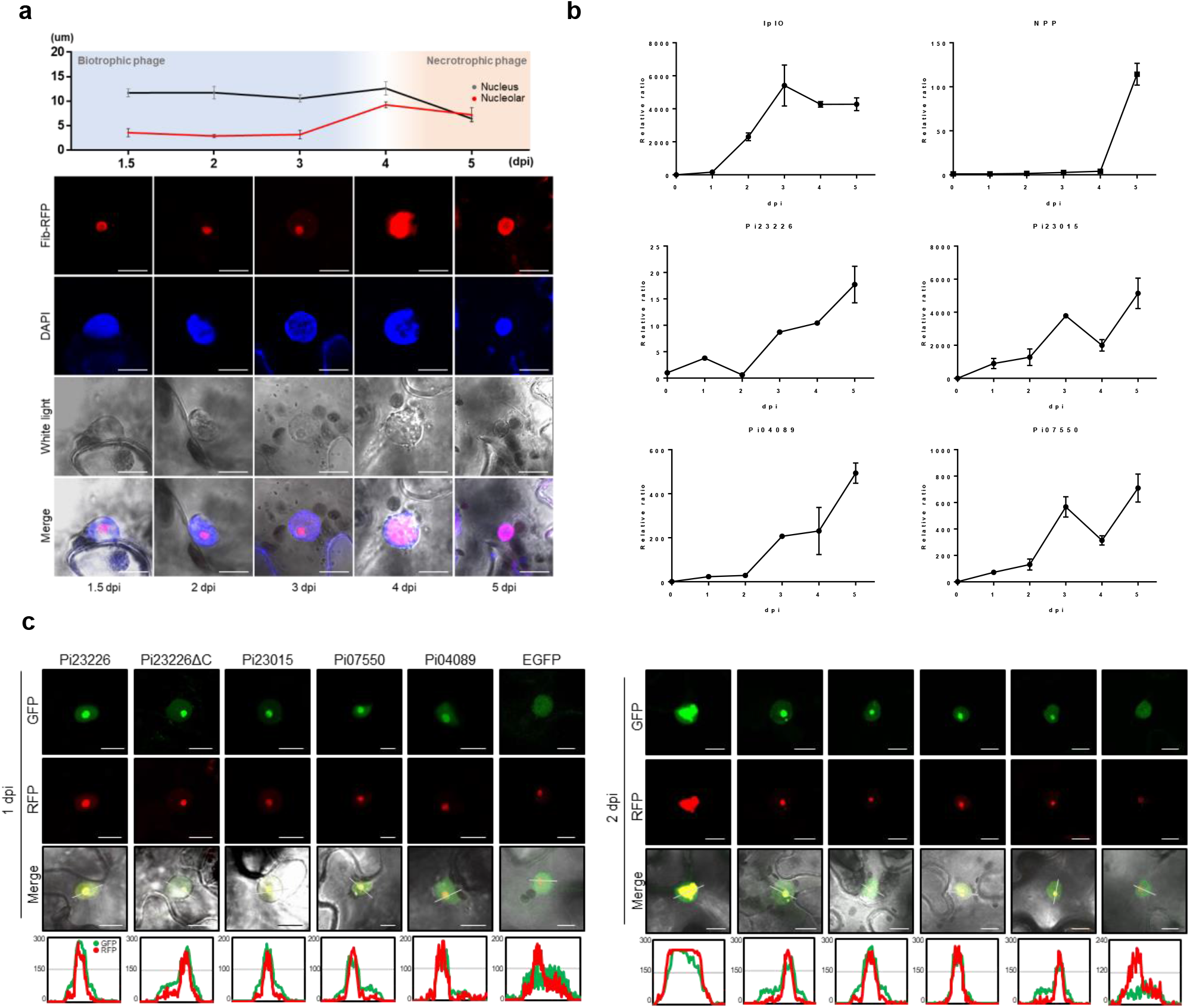
Inflation of host nucleoli before the initiation of necrosis following infection of *P. infestans*. (A) Morphology of host nucleolus during infection of *P. infestans*. RFP-fibrillarin2 (RFP-Fib2, red) was detected by confocal microscopy (SP8 X, Leica, Germany) as a nucleolar marker. DAPI staining (blue) was used to track nuclear and nucleolar sizes. Images were captured for 5 days after infection under red, blue, and white light, and subsequently merged. Nuclear and nucleolar sizes were measured. Representative confocal images from three independent experiments are presented. (B) Transcript levels of IpIO and NPP1.1, and nucleolar-localizing effectors were measured by qRT-PCR. Each transcript level was normalized to each gene at the initiation of infection (0 dpi). Representative fold changes are depicted as the mean values ± SD from three biological replicates. (C) Nucleolar localization of *P. infestans* effectors. Each GFP-fused effector was detected by confocal microscopy along with RFP-Fib2 as a nucleolar marker. Images were captured at 1 and 2 dpi under green, red, and white light, and subsequently merged. GFP signals from each effector construct were overlaid with those of RFP-Fib2 in the histogram. Representative confocal images from 10 independent experiments are presented.

We investigated the impact of pathogen infection on *N. benthamiana* nucleolus morphology using RFP-tagged fibrillarin2 (RFP-Fib2) as a nucleolar marker. No significant change in the nuclear or nucleolar size was observed during the biotrophic stage (ca. 1-3 dpi), whereas nucleolus inflation occurred at 4 dpi without a change of nuclear size (Figure 1A). Nuclear shrinkage and a reduction in nucleolar size occurred along with the progression of necrotic cell death. These results suggest that the pathogen modulates the host nucleolus activities during the switch of the disease cycle from the biotrophic to necrotrophic stage.

Several pathogen effectors have been reported to localize in the nucleolus, including Pi23226, Pi23015, Pi07550, and Pi04089 (Wang et al., 2019). Therefore, we transiently expressed these nucleolar-localized effectors in *N. benthamiana* to identify effector with similar functions in the plant nucleolus. The subcellular localization of the four effectors (Pi23226, Pi23015, Pi07550, and Pi04089) was confirmed by transient coexpression of C-terminal GFP-tagged effector constructs along with RFP-Fib2 as nucleolar marker in *N. benthamiana* leaves. All of the effectors colocalized with RFP-Fib2 primarily in the nucleolus (Figure 1C). Pi23015 and Pi23226 have almost identical amino acid sequences except for the C-terminal variance (Figure S1); therefore, we generated a deletion mutant containing only the conserved sequence (Pi23226ΔC). We observed that Pi23226ΔC was localized in the nucleolus, similar to Pi23226 and Pi23015, indicating that Pi23226ΔC still contains the nucleolar localization signal.

We examined the nucleolar morphology in the effector-expressing *N. benthamiana* cells. Nucleolar size inflation was observed only in Pi23226-expressing cells (Figure 1C). Pi23226 was reported to induce cell death in *N. benthamiana* (Lee et al., 2018), and we postulated that Pi23226 may induce cell death by modulating host nucleolar activity to enhance pathogen virulence. We investigated the impact of these nucleolar-localizing effectors during infection by measuring their transcript levels using qRT-PCR. The transcripts of Pi04089, Pi07550, Pi23015, and Pi23226 gradually increased during the biotrophic stage and reached maximum levels at 4 to 5 dpi (Figure 1B), which is considered as the time for transition to the necrotrophic phase. These results suggested that Pi23226 may function in the host nucleolus for the transition of pathogen life cycle from the biotrophic to necrotrophic phase.

### Nucleolar inflation coincides with Pi23226-induced cell death

Pi23226 induces host cell death concurrently with nucleolar size inflation. We tested other nucleolar effectors for their cell death-inducing activities. However, none of the other transiently expressed nucleolar effectors induced cell death in *N. benthamiana* leaves (Figure 2A). The inflation of nucleolar size observed in Pi23226-expressing cells occurred before visible cell death. We investigated whether nucleolar size inflation was resulted from the cell death phenotype by observing nucleolar morphology in cells expressing INF1 and CC309. INF1 and CC309 induce cell death by plasma membrane-mediated signal transduction in *N. benthamiana* (Lee et al., 2022). No change in nucleolar size was observed during cell death in cells expressing INF1, or CC309; however, nuclei were observed to shrink in all the cell death caused by expression of INF1, CC309 or Pi23226 (Fig 2b and c). We concluded that nucleolar size inflation coincided with cell death induced by Pi23226. Because nucleoli primarily function in ribosome biogenesis, we postulated that Pi23226 may affect ribosome biogenesis and protein translation, thereby leading to cell death.

**Figure 2.**
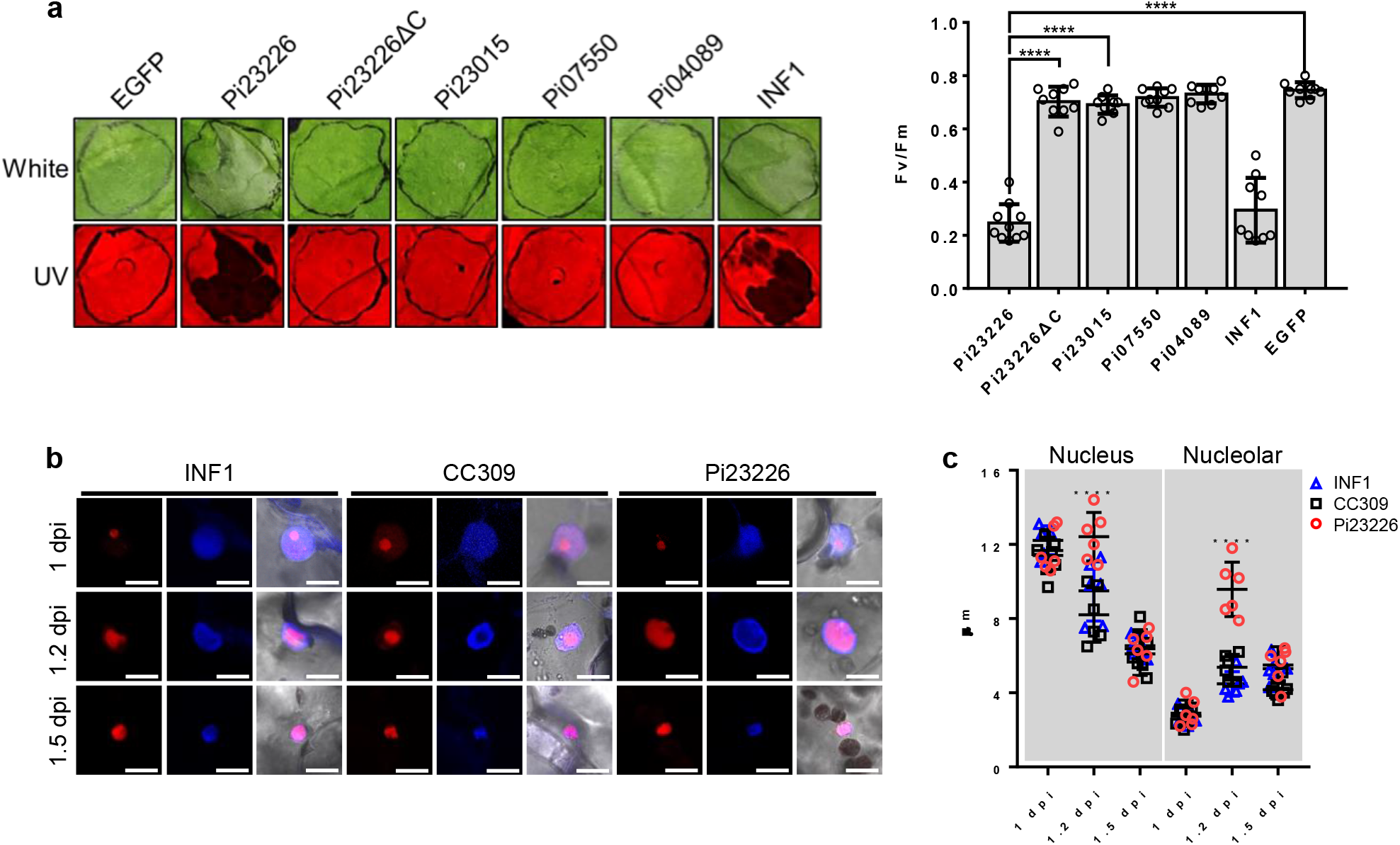
Nucleolar inflation coincides with Pi23226-induced cell death. (A) Cell death induced by the Pi23226 nucleolar effector. Cell death was observed under white and UV light. INF1 and mCherry were used as positive and negative controls, respectively. Representative images are presented. The quantum yield of photosystem II (Fv/Fm) was measured (Closed FluorCam FC 800-1010GFP, Photon Systems Instruments, Czechia) to represent the intensity of cell death. Data are mean values ± SD of five biological replicates. Significance was determined using one-way ANOVA (**** p<0.0001). (B) Nucleolar morphology in cells expressing Pi23226, INF1, and CC309. Each construct was coinfiltrated with RFP-Fib2 into *N. benthamiana*, and the nucleus and nucleolus were observed during the progression of cell death. Images were captured for two days after infection under red, blue, and white light, and subsequently merged. Representative confocal images from three independent experiments are presented. (C) Changes in nuclear and nucleolar sizes in cells expression Pi23226, INF1 or CC309. Each Pi23226, INF1 or CC309 construct was coinfiltrated with RFP-Fib2 into *N. benthamiana* leaves. Nuclear and nucleolar diameters were measured using ImageJ at 1, 1.2, and 1.5 dpi. For oval shapes, the semi-major axis was measured. Data are mean values ± SD for three biological replicates. Significance was determined using one-way ANOVA (**** p<0.0001).

### Pi23226 directly binds 25S rRNA

To test whether nucleolar inflation is caused by disrupted ribosome biogenesis, we examined the binding of Pi23226 to rRNA-RNP complexes using RNA immunoprecipitation (RIP) of transiently expressed FLAG-tagged effectors in *N. benthamiana*. RNA-protein complexes were crosslinked with formaldehyde, precipitated with anti-FLAG affinity gels, and followed by reverse crosslinking. The amount of precipitated RNA was measured by qRT-PCR analysis using primers amplifying 18S, 5.8S, and 25S rRNA and normalized to the input, which was similarly treated as RIP but without precipitation. The 25S rRNA region was amplified 20- to 30-fold in cells expressing Pi23226, whereas no amplification of the 25S rRNA region was observed in cells expressing Pi23226ΔC or Pi23015 (Figure 3A and B). By contrast, the 18S and 5.8S regions were not enriched in any of the samples. To investigate the direct interaction of Pi23226 with rRNA, we performed individual-nucleotide resolution UV crosslinking and immunoprecipitation (iCLIP), followed by next-generation sequencing (NGS) to precisely identify the Pi23226-binding sequence of 25S rRNA. Consistent with the RIP results, iCLIP-NGS analyses identified the sequence covering the 3′-end of the 25S rRNA gene as the Pi23226-binding region (Figure 3C and S2). To verify the NGS results, we designed primer sets to amplify the exact 3′-end region identified in the NGS results (primer 25S-4) and covering the peripheral area (primer 25S-3 and 25S-5) (Figure S3). The qRT-PCR results showed that the 25S-3 amplicon was enriched in Pi23226-expressing cells but not in Pi23226ΔC- or Pi23015-expressing cells (Figure 3D). Amplicons at the peripheral area (primers 25S-3 and 25S-5) were not enriched in any of the construct-expressing samples. These results indicate that Pi23226 directly binds to rRNA, especially the 3′-end of the 25S rRNA region (Figure 3A). Thus, we postulated that Pi23226 binding to the 25S rRNA sequence may disrupt 25S rRNA processing from its precursors.

**Figure 3.**
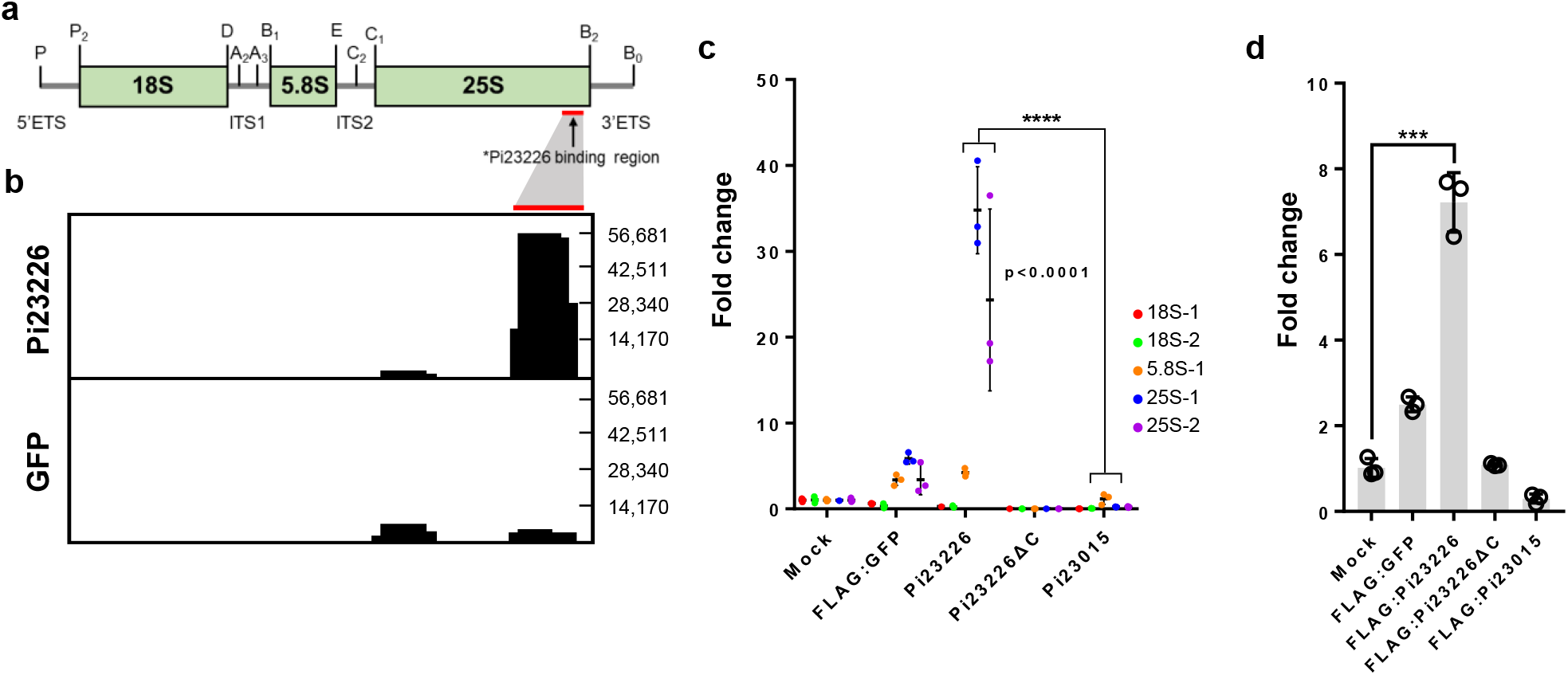
Pi23226 directly binds to 25S rRNA. (A) 45S rRNA transcript containing 18S, 5.8S, and 25S rRNAs. Exonucleolytic and endonucleolytic cleavage sites are indicated. The Pi23226 binding region on 25S rRNA from iCLIP is marked with a red bar. (B) RIP using FLAG-fused effector constructs. FLAG-effector constructs were expressed in *N. benthamiana* using agroinfiltration. The abundance of RNAs associated with FLAG-fused effectors was measured and normalized relative to inputs which were processed identically as RIP except for precipitation. Specific primers to amplify 18S, 5.8S, and 25S rRNAs were used in qRT-PCR. Data represents mean values ±SD, and significance was determined using Student’s t-test (**** p<0.0001). (C) Pi23226 specifically binds to the 3′-end of 25S rRNA. iCLIP-seq data were mapped on the *N. benthamiana* genome (v1.0.1). The normalized mapped reads on 25S rRNA gene region are presented as Integrative Genomics Viewer. iCLIP-seq data from FLAG:GFP-expressing *N. benthamiana* are used as a negative control. (D) iCLIP-qRT-PCR for validation of iCLIP-seq. FLAG-effector constructs were expressed in *N. benthamiana*. iCLIP-qRT-PCR was performed using primers derived from the 3′-region of 25S rRNA (Figure 2b). RNA abundance was measured and normalized relative to inputs, which were processed identically as iCLIP except for precipitation. Representative fold changes of amplicons are depicted as mean values ±SD from five biological replicates and significance was determined using Student’s t-test (*** p<0.001)

### Pi23226 disrupts pre-rRNA processing and induces nucleolar stress

To investigate whether Pi23226 binding to 25S rRNA affects rRNA processing, we measured the accumulated levels of mature rRNAs and their byproducts using qRT-PCR. Unexpectedly, all 18S, 5.8S, and 25S rRNA fragments were highly accumulated in cells expressing Pi23226, Pi23226ΔC, and Pi23015 compared to the EGFP control, whereas the 5′-ETS, ITS1, and ITS2 regions did not show a significant difference. There was no abnormal accumulation of rRNA in cells expressing INF1 compared to the EGFP control, indicating that rRNA accumulation was not a result of cell death (Figure 4A). This suggests that Pi23015 and Pi23226ΔC may increase mature rRNA accumulation by an unknown mechanism. A 10-fold higher accumulation of 3′-ETS was observed only in cell expressing Pi23226 (Figure 4A). The ratio of untrimmed 25S rRNA precursor/25S rRNA also increased only in cells expressing Pi23226 (Figure 4B), suggesting that Pi23226 binding to the 3′-end of the 25S rRNA precursor effectively disrupted cleavage of the 3′-ETS from the 25S rRNA precursor. To confirm that Pi23226 disrupted pre-rRNA maturation, we performed circular RT-PCR (cRT-PCR). Each fragment of 18S, 5.8S, and 25S was circularized by reverse primers, designed as 18C, 5.8C, and 25C, respectively, subjected to PCR with the indicated primer sets, and detected in agarose gels (Figure 4C-1). Two abnormal bands were detected only for 25S cRT-PCR in cells expressing Pi23226, whereas no significant differences were observed in cells expressing other effector constructs (Figure 4C-2). Sequencing analysis of the abnormal bands indicated that one contained 27SB including 5.8S, ITS2, and 25S rRNA sequences, whereas the other contained 27SA_3_ and part of 3′-ETS (3′-ETS_p_). This indicated that ectopic expression of Pi23226 caused inappropriate processing in the maturation steps from 27SA_3_ and 27SB rRNA to 25S rRNA precursors and the removal step of 3′-ETS from 25S rRNA precursors. This also suggests that Pi23226 binding on the 3′-end of 25S may disrupt the early stage of the ITS1-first pathway in pre-rRNA processing and result in the accumulation of malprocessed pre-rRNA.

**Figure 4.**
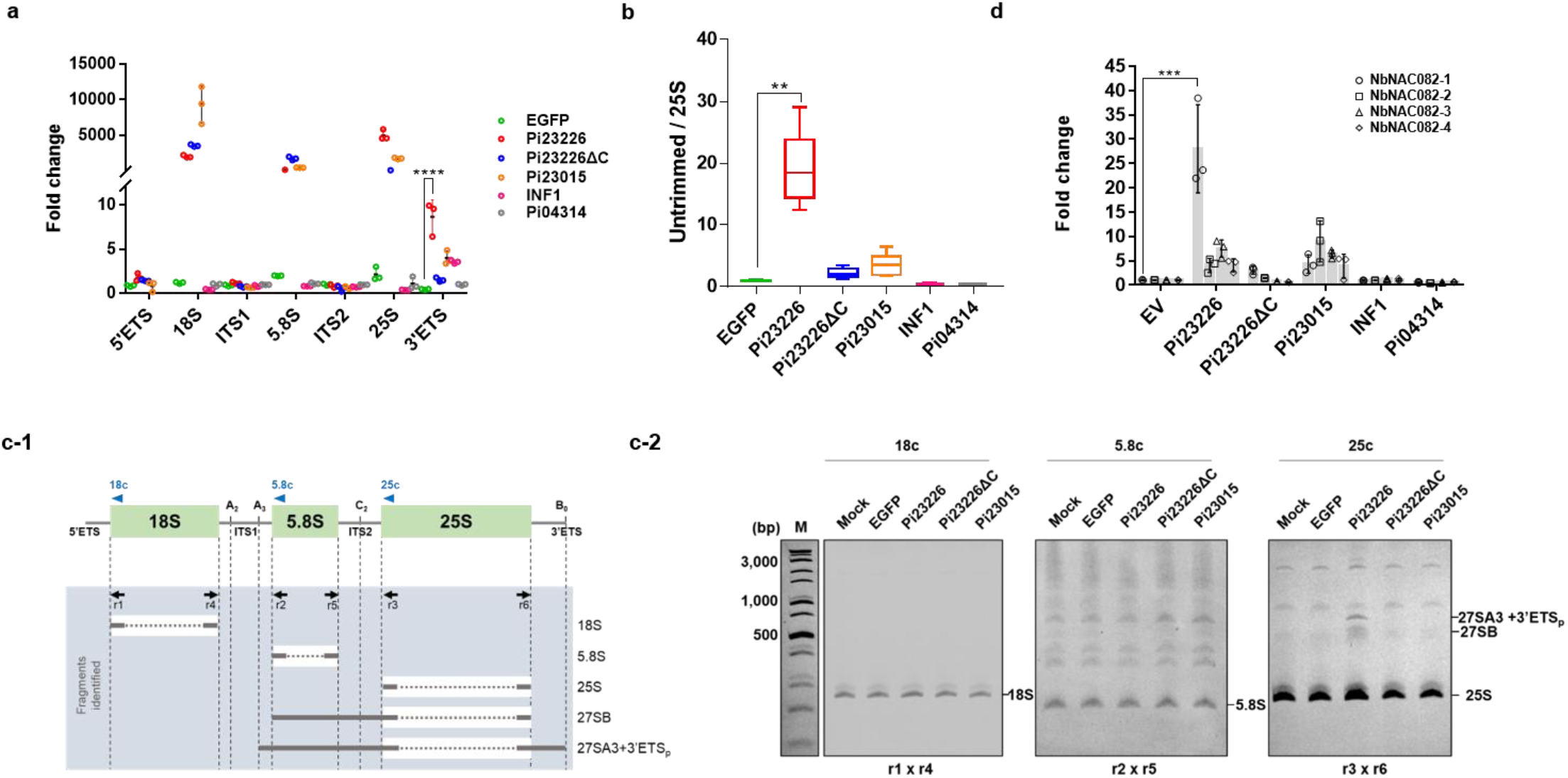
Pi23226 disrupts pre-rRNA processing and induces nucleolar stress. (A) FLAG-fused effector construct and INF1 expression change the rRNA fragment repertoires. The abundance of pre-rRNA fragments was measured after the expression of Pi23226, Pi23226ΔC, Pi23015, and INF1 using qRT-PCR. The graph depicts the relative ratios of amplified fragments normalized to the level of ubiquitin (UBQ) transcripts. RNA samples from eGFP-expressing *N. benthamiana* cells were used as a negative control. A representative graph from four biological replicates is presented, and the significance was determined using one-way ANOVA (**** p<0.0001). (B) Pi23226-induced accumulation of unprocessed 27S rRNAs. The untrimmed 3′-region of 25S rRNA was quantified by qRT-PCR using primers covering 25S and 3′-ETS. The relative ratio was calculated from the amplicons covered from the end of 25S rRNA to 3′-ETS normalized by the level of 25S rRNA transcripts. RNA samples from eGFP-expressing *N. benthamiana* cells were used as a negative control. A representative graph from six biological replicates is presented, and the significance was determined using one-way ANOVA (*** p<0.001). (C-1) Primers designed for cRT-PCR to amplify the pre-rRNA fragments. Nuclear RNA samples from effector-expressing *N. benthamiana* cells were circularized by RNA ligation; 18S, 5.8S, and 25S intermediate cDNAs were synthesized using specific primers (arrows without bars); and each intermediate was amplified by PCR (arrows with bars). Amplicons in Figure 3C-2 are depicted as black bars. (C-2) Gels showing the rRNA processing intermediates using cRT-PCR with different primer sets as indicated in Figure 3C-1 under effector-expressing conditions. Bands at the bottom present the minimal size, which can be amplified with each primer set following cRT-PCR. Abnormal bands shown in the cRT-PCR by 25c under Pi23226-expressing conditions represent 27SA3 and 27SB. Representative images from three biological replicates are shown. (D) Transcript levels of NAC082 transcription factor candidates measured by qRT-PCR. Each transcript level was normalized to EF-α as a reference gene and compared to eGFP as a negative control. Significance was determined using one-way ANOVA (*** p<0.001).

These observations of Pi23226-induced enlargement of nucleolar size and accumulation of malprocessed rRNA prompted us to investigate the activation of nucleolar stress. The p53 transcription factor is a key regulator of the ribosomal stress response in mammals, although no p53 ortholog has been reported in plants. The Arabidopsis NAC082 transcription factor (ANAC082, AT5G09330) has been reported as an essential biomarker that is upregulated under nucleolar stress conditions (Ohbayashi et al., 2017). Therefore, we examined the transcript levels of the NAC082 transcription factor by qRT-PCR. We selected four NAC082 transcription factor candidates from *N. benthamiana* (designated as NbNAC082-1 to −4 according to the degree of similarity to ANAC082). *NbNAC082-1*, the closest homolog of ANAC082, was specifically upregulated only in Pi23226-overexpressing cells (Figure 4D). These results indicate that Pi23226 binding to the 3′-end of the 25S rRNA sequence may lead to the accumulation of untrimmed 25S rRNA by inappropriate processing from 27S to 25S rRNA, which eventually affects ribosome biogenesis and causes nucleolar stress.

### Pi23226 disrupts protein synthesis by suppressing global translation

We investigated whether Pi23226-induced defects in pre-rRNA processing perturb ribosome assembly by generating a polysome profile using ultra-High Performance Liquid Chromatography(uHPLC). Polysomes were isolated from leaf tissues expressing FLAG:Pi23226, FLAG:dCPi23226, or FLAG:Pi23015, and then fractionated into 27 columns with a column flow rate of 0.4 ml/min with monitoring absorbance at 260 nm (Figure 5A). Polysomes isolated from untreated and FLAG:GST-expressing leaf tissues were used as negative controls. To validate the relevant protein distribution in polysome profiling, proteins were extracted from each fractionated column and immunoblotted using a ribosomal protein large subunit 13 antibody (α-RPL13). Pi23226 caused a significant increase in 60S and reduced the 80S/60S ratio, whereas Pi23226ΔC and Pi23015 did not affect the distribution of ribosomes (Figure S4). To investigate whether Pi23226 associated with ribosome subunits, we performed western blotting using α-FLAG. The results showed that Pi23226, but not Pi23226ΔC or Pi23015, was detected in fractions representing polysomes and 60S subunits. We postulated that Pi23226 association with polysomes and rRNA may affect pre-rRNA processing and assembly; thus, we investigated the effect of Pi23226 on protein translation activity. First, we adapted the wheat germ extract system for *in vitro* translation experiments. Recombinant *GFP* plasmid DNA was used as a template for translation. We validated the system by inhibiting the *in vitro* translation of GFP using the translation inhibitor cycloheximide (CHX). The translated GFP products were significantly reduced according to the CHX concentration (Figure 5B, upper panel). Subsequently, the recombinant GST:Pi23226, GST:Pi23226ΔC, and GST:Pi23015 proteins were incubated with wheat germ extract in the presence of *GFP* plasmid constructs (Figure S5). GFP protein was not translated in the presence of GST:Pi23226, whereas neither GST:Pi23226ΔC nor GST:Pi23015 affected the translation of GFP (Figure 5B, lower panel). Next, we examined *de novo* protein synthesis *in planta* using a surface sensing of translation (SUnSET) assay (Van Hoewyk, 2016). Similar to the *in vitro* translation system, *de novo* protein was barely detected by treatment with a high concentration of CHX (Figure 5C, left panel). Consistent with the *in vitro* analysis, *de novo* protein translation was significantly reduced under Pi23226-expressing conditions, whereas no significant change in total protein synthesis was observed under Pi23226ΔC or Pi23015-exxpressing conditions (Figure 5C, right panel). These combined results suggest that Pi23226 but not Pi23226ΔC or Pi23015 disrupts protein synthesis, probably by associating with functional polysomes and the 60S subunit, thereby resulting in translation inhibition.

**Figure 5.**
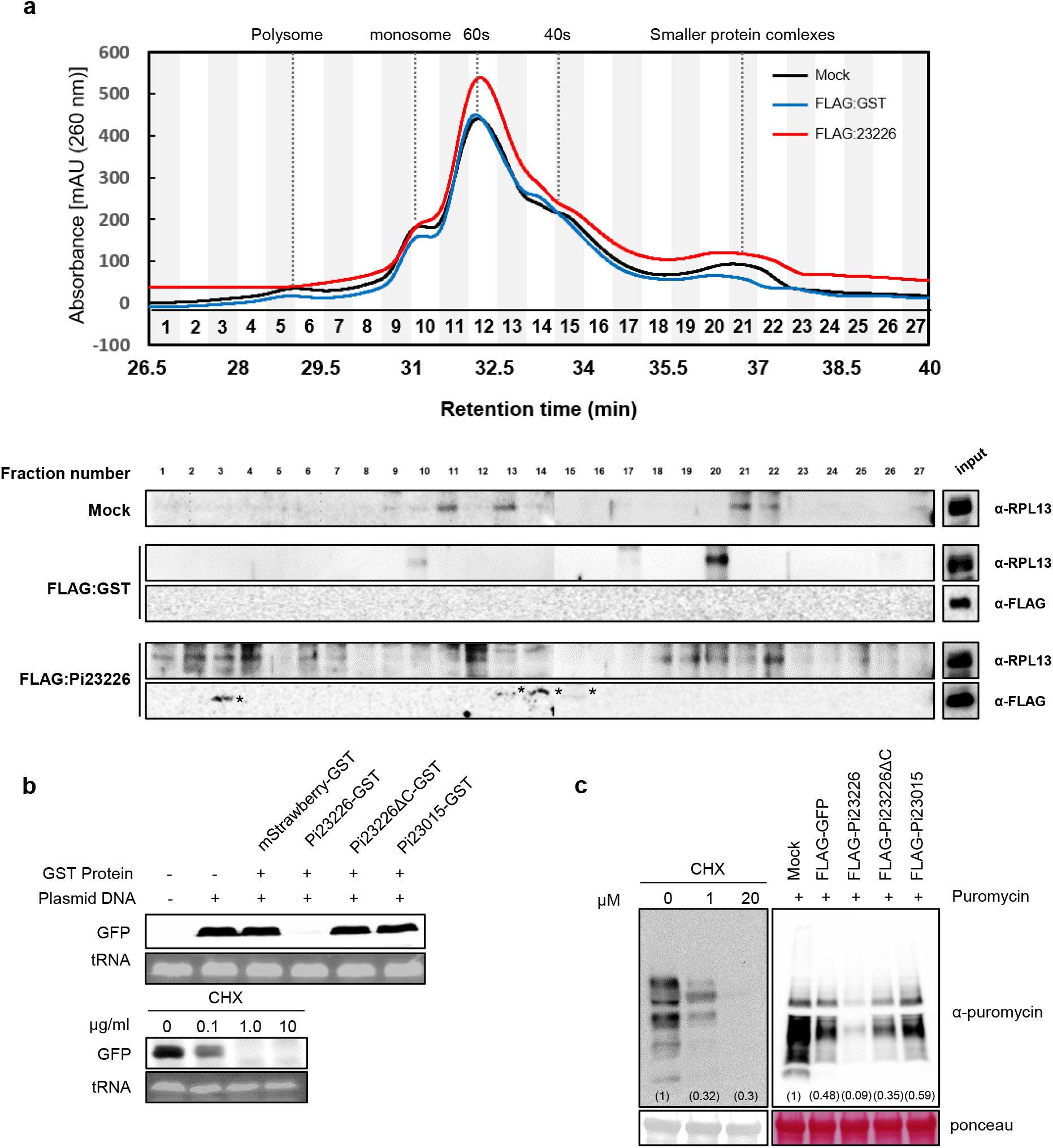
Pi23226-induced interruption of protein synthesis suppresses global translation. (A) Pi23226 affected the polysome profile. Ribosomes were isolated from leaf tissues expressing FLAG:Pi23226. Untreated or FLAG:GST-expressing conditions were used as negative controls. Fractions were collected into 27 columns with a column flow rate of 0.4 ml/min with monitoring absorbance at 260 nm. Fractionated proteins were extracted and detected with an anti-RPL13 or anti-FLAG antibody and compared with input. Representative images from three biological replicates are shown. (B) Pi23226 inhibits *in vitro* translation of *GFP* using wheat germ extract. Left panel: CHX-dependent inhibition of *in vitro* GFP translation. Right panel: GFP translation *in vitro* following incubation with GST-fused recombinant effector proteins. GFP was detected with an anti-GFP antibody. Equal loading was controlled by adding ε-labeled biotinylated lysine tRNA. Representative images from five independent experiments are shown. (C) Pi23226 inhibits *de novo* protein synthesis in the SUnSET assay. The level of newly synthesized protein from effector-expressing *N. benthamiana* following puromycin treatment was measured using an anti-puromycin antibody. Left panel: CHX-dependent inhibition of *de novo* protein synthesis. Protein samples were extracted after puromycin treatment following CHX supplementation. Right panel: SUnSET assay for the detection of newly synthesized protein in FLAG-effector-expressing *N. benthamiana*. FLAG:GFP-expressing and untreated leaf tissues were used as negative controls. The loading control was visualized by Ponceau staining.

## Discussion

In the hemibiotrophic life cycle of pathogen infection, cell death can be harmful or beneficial to pathogens depending on the timing of cell death. Plant immune receptors can recognize effector activities and induce HR cell death as an immune response. Most of the disease-resistant HR cell death induced by immune receptor recognition is associated with the plasma membrane and cytosol (Bi et al., 2021; Lee et al., 2020, 2021). Pathogens also facilitate host cell death to gain benefits from the dying tissue by transiting from the biotrophic to the necrotrophic phase. Thus, they utilize cell death by hijacking essential components of the host cell in diverse subcellular organelles without immune receptor recognition. However, it is not clear when and how pathogens shift their lifestyle from the biotrophic to the necrotrophic phase. In this study, we observed nucleolar inflation concurrently with the switch from biotrophic to necrotrophic stage during *P. infestans* infection. This suggests that cell death signaling initiated from the nucleolus could be a strategy mediating the transitioning of the disease cycle. The nucleolar-localized Pi23226 induced a similar nucleolar morphology as that induced by the disease cycle transition when expressed in *N. benthamiana*. Pi23226 binding to the 25S rRNA disrupted ribosome biogenesis, reduced proteins translation, and may trigger necrotic cell death of the host, resulting in enhanced host susceptibility to the pathogen (Figure S6). We also observed that Pi23226 suppressed global protein translation *in vitro* and *in planta*, suggesting that Pi23226 may hijack ribosomes and induce nucleolar stress responses leading to host cell death. Our results and previous studies suggested that *P. infestans* may utilize nucleolar-localizing effectors to induce the disease transition in host cells.

As genome sequences of major plant pathogens become available, hundreds of pathogen effectors have been identified and their functions in host cells have been reported (Gao et al., 2020; He et al., 2020). Most of the effector targets are host proteins, and only a few effectors are localized in nuclei and directly bind to host DNA, such as the transcription activator-like effectors (TALEs) (Kay et al., 2007; Szurek et al., 2001). A small number of effectors from bacterial and filamentous pathogens are localized in the nucleus and nucleolus (Boevink et al., 2016; Kim et al., 2020), although none have been confirmed to directly bind RNA. A recent study tried to identify RNA-binding effectors from a collection of animal and plant pathogen effectors; however, only a small number of effectors were reported that were theoretically predicted to carry RNA binding domains (Tawk et al., 2017). To the best of our knowledge, our results provide the first evidence of pathogen effectors that directly bind to RNA *in vivo*. The current study demonstrated that the *P. infestans* effector Pi23226 directly associated with rRNA, especially at the 3′-end of 25S rRNA, disrupts ribosome biogenesis, and inhibits translation to enhance pathogenicity. The binding of Pi23226 to 25S rRNA precursors appeared to interfere with the processing of 27S pre-rRNA to 25S rRNA because the quantity of untrimmed 3′-ETS increased in cells expressing Pi23226, suggesting that endoribonucleolytic cleavage of the B2 site at the 3′-end of 25S rRNA was incomplete. It is unlikely that Pi23226 had endonuclease activity, as it did not contain any domain identified in RNase. Pi23226 is a relatively small protein (123 amino acids) with no known functional motif, and appears to act as a small interfering adaptor. Thus, Pi23226 binding on the end of 25S rRNA precursors may block the access of RNase to the cleavage site in the 3′-ETS. A ribonuclease involved in cleavage of the 3′-ETS in *N. benthamiana* has not been identified, but RNASE THREE LIKE2 (RTL2) requires cleavage of the pre-rRNA in the 3′-ETS in Arabidopsis (Comella et al., 2008). Therefore, it would be interesting to further investigate whether Pi23226 binding on the 25S rRNA precursors affects the activity of RTL2 or a homolog on the cleavage of 3′-ETS in *N. benthamiana*. The accumulation of the 27SA_3_+3′-ETS and 27SB pre-rRNAs under Pi23226-expressing conditions suggests that the increase in untrimmed 3′-ETS could suppress further processing of 27S rRNA to 25S rRNA, which is recognized by the ribosome biogenesis surveillance system and induce nucleolar stress responses leading to cell death.

Although we showed that Pi23226 had the RNA-binding activity, we do not rule out the possibility that Pi23226 also associated and functioned with yet-unidentified host proteins. The polysome profiling analyses detected Pi23226 on the polysome fraction (fraction 3, Figure 4A), suggesting that Pi23226 may function on polysomes by associating with rRNA or other proteins, and we previously reported that Pi23226 is associated with heat shock protein 70 (HSP70) (Lee et al., 2018). The Pi23015 homolog of Pi23226 was reported to have a role in alternative splicing of pre-mRNAs and to promote pathogenicity (Huang et al., 2020).

In conclusion, our results provide the first *in vivo* evidence of RNA-binding pathogen effectors, and extends our knowledge of the nucleolar action of pathogen effectors that disrupt ribosome biogenesis.

## Materials and Methods

### Plant growth conditions and Agrobacterium-mediated transient overexpression in plants

Plants were grown and maintained in a work-in chamber with an ambient temperature of 23– 24°C and 16 h/8 h (day/night) cycle. For *in planta* expression, *A. tumefaciens*-mediated transient expression was performed on 4-week-old *N. benthamiana* plants. An overnight culture of *A. tumefaciens* strain GV3101 was centrifuged, resuspended in buffer (10 mM MgCl_2_, 10 mM MES-KOH pH 5.7, 200 μM acetosyringone) and incubated for 1 h at room temperature. The bacterial suspension was adjusted to OD600 = 0.4 and pressure-infiltrated into the abaxial surface of *N. benthamiana* leaves using a needleless syringe. For coexpression experiments, Agrobacteria harboring each construct were mixed and adjusted to OD600 = 0.4 in the same suspension and infiltrated. Leaf tissues were collected at 1.5∼2 dpi.

### Plasmid constructs

For N-terminal GFP fusions, RXLR effector genes (Pi23226, Pi23015, Pi07550, and Pi04089) and other mutants were synthesized without a signal peptide and RXLR regions, were amplified with primers containing the NruI restriction site, and then ligated into the PZP212 vector using enzyme digestion. Fibrillarin was amplified with the attB site from cDNA of *N. benthamiana* and fused to RFP using pB7WGF2 in the GATEWAY system. To detect the protein level of effectors, the amplicons of 3×FLAG-Pi23226 and 3×FLAG-mutants were cloned into the pkw-LIC vector containing the cauliflower mosaic virus 35S promoter using the ligation-independent cloning (LIC) method (Oh et al., 2010). Generated vector constructs were transformed into *A. tumefaciens* GV3101 for transient expression. The primer sequences used in this study are presented in Table S1.

### Confocal microscopy

To determine the subcellular localization of GFP-tagged Pi23226 in *N. benthamiana* leaves, samples were observed using Leica SP8 X (USA) confocal microscopes equipped with a ×40 water immersion objective. GFP fluorescence was visualized using a 488 nm laser line for excitation, and emission was collected between 455 and 480 nm. For imaging mRFP fluorescence, excitation was at 532 nm, and emission was observed between 580 and 650 nm. To minimize spectral bleed-through between fluorescence channels during colocalization experiments, images were acquired by sequential scanning with alternation between frames. Plant nuclei were stained using DAPI (4′,6-diamidino-2-phenylindole, 10 μg/ml in water) that was pressure-infiltrated into the abaxial side of the leaf 10 min before observation. DAPI fluorescence was excited at 405 nm and observed between 450 and 480 nm. Nuclear size was measured and analyzed using ImageJ software.

### Protein isolation and immunoblot analyses

Total protein was extracted using extraction buffer [10% (v/v) glycerol, 25 mM Tris-HCl (pH 7.5), 1 mM EDTA, 150 mM NaCl, 1% (w/v) polyvinylpolypyrrolidone, and 1× protease inhibitor cocktail], mixed with SDS/PAGE sample buffer, and analyzed by immunoblotting. For western blotting, protein samples were separated on SDS-PAGE gels and transferred to PVDF membranes (Bio-Rad, 88585) using a Trans-Blot Turbo Transfer System (Bio-Rad, 1704150). Membranes were blocked with 6% milk prepared in Tris-buffered saline with 0.1% Tween 20 (TBST) and incubated with primary anti-FLAG (1:12000, Sigma-Aldrich, F3165) or anti-GFP (1:12000, Invitrogen, A10260) antibodies at room temperature for 1 h. The membrane was washed twice with PBST for 10 min each before the addition of the secondary anti-mouse Ig-HRP antibody (1: 15,000, Abcam, ab6708) or anti-rabbit Ig-HRP antibody (1:15,000, Abcam, ab6702) for 1 h. ECL (Bio-Rad, 1705061) detection was performed according to the manufacturer’s instructions.

### *In vitro* and *in vivo* translation assay

*In vitro* translation was carried out using a wheat germ protein expression system (Promega, USA) following the manufacturer’s instruction. In brief, 2 μg of GFP pF3WG, 1 μg of GST-tagged effector proteins and 2 μl of FluoroTect™ GreenLys tRNA (Promega, USA) were gently mixed with High-Yield Wheat Germ Master Mix, followed by incubation at 25°C for 2 h. Recombinant GST-strawberry was used as a negative control for comparison with other GST recombinant proteins. The final protein products were analyzed by SDS-PAGE and immunoblotting with anti-GFP, and the level of tRNA was detected using an Azure 400 chemiluminescence imager (Azure Biosystems, USA) as an equal loading control. For cycloheximide (CHX) treatment, CHX (Amresco, 94271) at the indicated concentrations was added together with Wheat Germ Master Mix before incubation. *De novo* protein synthesis was examined *in planta* using a modified surface sensing of translation (SUnSET) assat (Van Hoewyk, 2016). Briefly, *N. benthamiana* leaf discs were incubated with 100 μM puromycin for 2 h in 1/2 MS medium under vacuum. Then, 30 μg of total protein extract was loaded onto a 4–20% gradient SDS-PAGE gel, and puro-polypeptides were detected by western blotting with an anti-puromycin antibody (Millipore, MABE343) at a dilution of 1:10,000. Newly synthesized proteins were detected using an Azure 400 chemiluminescence imager (Azure Biosystems, USA), and the intensity of *de novo* protein synthesis was calculated using ImageJ. For CHX treatment, leaf discs were incubated with CHX in 1/2 MS medium at 25°C for 4 h before puromycin treatment.

### Polysome isolation and profiling

Ribo Mega-SEC was modified for polysome profiling (Yoshikawa et al., 2018). Approximately one-half of the *N. benthamiana* leaves were ground in liquid nitrogen, homogenized in 500 μl of polysome isolation buffer (200 mM Tris-HCl pH 8.5, 50 mM KCl, 25 mM MgCl_2_, 1% sodium deoxycholate, 1% CHAPS, 50 μg/ml chloramphenicol, 50 μg/ml cycloheximide, 1× protease inhibitor cocktail, 400 U/ml SUPERase In RNase Inhibitor), and incubated for 15 min on ice. Then, the homogenate was clarified by centrifugation at 17,000×g for 10 min at 4°C. The supernatant was passed through a 0.45 μm Ultrafree-MC-HV centrifugal filter (UFC30HVNB, Merck Millipore, USA) followed by centrifugation at 12,000×g for 2 min at 4°C. Then, 25 μg of ribosome isolate was injected with SEC buffer (20 mM Hepes-NaOH pH 7.4, 130 mM NaCl, 10 mM MgCl_2_, 1% CHAPS, 2.5 mM DTT, 50 mg/ml cycloheximide, 20 U SUPERase In RNase Inhibitor, cOmplete EDTA-free Protease Inhibitor Cocktail) and analyzed by chromatography system (UltiMate 3000 UHPLC, Thermo Fisher Scientific) using Agilent Bio SEC-5 2,000 Å columns (7.8 × 300 mm with 5-mm particles). The gradient was collected with simultaneous measurement of UV absorbance at 260 nm (flow rate 0.4 ml/min). Fractionated proteins were isolated with MetOH/chloroform and detected by western blotting with the 60S ribosomal protein L13-1 and antibody (Agrisera, AS132650, Sweden) at a dilution of 1:4,000.

### UV crosslinking and immunoprecipitation (iCLIP)-seq and RIP

Four-week-old *N. benthamiana* leaves were irradiated three times with UV at 0.500 J/cm^2,^ and leaf samples were ground in liquid nitrogen. Total protein extracts were filtered through a 0.45 µm filter and sonicated in an ice bath using a bioruptor (Sonifier SFX 550, USA) at 30-s on/30-s off for 15 min. Samples were centrifuged at 16,000 × g for 20 min, and the supernatant was collected. Supernatants were treated with 4 µL of DNAse I (NEB, EN0521) and incubated for 10 min at 37°C with shaking at 1,200 rpm. For RNA digestion, 40 U of RNase I was added for 10 min at 37°C with shaking at 1,200 rpm. The sample was precleared with agarose beads in IP buffer [10% (v/v) glycerol, 25 mM Tris–HCl pH 7.5, 1 mM EDTA, 150 mM NaCl, 1× protease inhibitor cocktail] for 1 h at 4°C. The sample was centrifuged, and the supernatant was incubated with a-FLAG-affinity agarose beads (BioLegend, USA) for 2 h at 4°C. The beads were washed four times with IP buffer followed by elution with 130 µl of elution buffer (1% SDS, 0.1 M NaHCO_3_, 5 units of SUPERase In RNase Inhibitor) by rotating for 30 min at 4°C. For reverse crosslinking, 10 μl proteinase K (20 mg/ml) was added and incubated at 60°C for 1 h. RNA was recovered with TRIzol reagent according to the manufacturer’s instructions. The genome sequences and annotations for genes and repeats in *N. benthamiana* were downloaded from the Sol Genomics Network (https://solgenomics.net/ftp/genomes/Nicotiana_benthamiana/) (Bombarely et al., 2012). The raw reads from iCLIP-seq were trimmed to remove sequences with low quality and adaptors using fastp v 0.20.1 (-l 71 -q 30) (Chen et al., 2018). The filtered reads were mapped to the reference genome in *N. benthamiana* using Hisat2 v 2.1.0 with the default parameter (Kim et al., 2019). The BAM files from aligned reads were sorted using Samtools v 1.10 (Li et al., 2009) and converted into raw and RPKM-normalized bed files using bamCoverage implemented in Deeptools v 3.5.0 (Ramírez et al., 2016). The raw and RPKM-normalized bed files were visualized in IGV v 2.8.4 (Robinson et al., 2011).

### Circular RT-PCR analysis

Circular RT-PCR analysis was performed using 10 µg of RNA extracted from nuclei of *N. benthamiana* leaves. For this, 1g of frozen leaves was suspended in nuclear isolation buffer (10 mM Tris-HCl, pH 8.0, 10 mM MgCl_2_, 400 mM sucrose, 10 mM β-mercaptoethanol, 0.2 mM PMSF, and 1× protease inhibitor cocktail). The solution was filtered using a cell strainer and centrifuged at 4°C for 20 minutes at 1,900×g. The pellet was resuspended in storage buffer [10 mM Tris-HCl, pH 8.0, 10 mM MgCl_2_, 250 mM sucrose, 0.15% (w/v) Triton X-100, 5 mM β-mercaptoethanol, 0.1 mM PMSF, and 1× protease inhibitor cocktail]. The quality and quantity of nuclei were confirmed under a microscope. RNA was extracted using TRIzol reagent (MRC, USA) and ligated with T4 RNA ligase 1 (NEB, USA) at 37 °C for 1 h. First-strand cDNA was reverse-transcribed from ligated RNAs using SuperScript III reverse transcriptase (Invitrogen, USA) and primers 18c, 5.8c, and 25c. The rRNA intermediates were amplified with specific primer sets (Table S1), and amplified PCR fragments were determined by sequencing.

### RNA extraction and gene expression analyses

Total RNA was extracted from *N. benthamiana* leaves using TRIzol reagent (MRC, USA). To investigate the level of rDNA transcription, samples expressing each construct were collected before the cell death phenotype was observed using the same amounts of leaf tissues. The same volumes of RNA were loaded on 1.5% agarose gel.

For gene expression analysis, reverse transcription was performed using SuperScript III reverse transcriptase (Invitrogen, USA) and random hexamer priming. Quantitative RT-PCR assays were conducted using PowerUP SYBR green master mix (Thermo, USA) according to the manufacturer’s instructions. The expression levels of genes were calculated using the 2-ΔΔCt algorithm (KJ and TD, 2001). Primer pairs were designed for each of the selected genes (Table S1). The *N. benthamiana* elongation factor 1α (*NbEF1α*) gene was used as an internal control.

### *Phytophthora* infection assays

The virulence function of effectors was evaluated using *A. tumefaciens* infiltration of *N. benthamiana* leaves. Each GFP-fused effector was transiently overexpressed on one-half of the leaf, and GFP was overexpressed on the other half of the same 4-week-old leaf. *P. infestans* isolate T30-4 was cultured on Rye agar at 19°C for 10 days. Plates were flooded with 5 ml cold H_2_O and scraped with a cell scraper to release sporangia. The zoospore solution was collected and diluted to 50,000 spores/ml. Infection assays were performed using droplet inoculations (10 μl of a zoospore solution) on detached leaves after 24 hpi. The lesion size was measured 5 days after inoculation. Each lesion size was photographed using Cy5 and Cy3 channels in an Azure 400 (Azure Biosystems, USA). and autofluorescent areas were analyzed using ImageJ software.

### Detection of cell death intensity

Cell death intensity was quantified by measuring chlorophyll fluorescence. Detached *N. benthamiana* leaves were exposed to a measuring beam, and the minimal (F_v_) and maximal (F_m_) fluorescence were determined using FluorCam 800MF (Photon Systems Instruments, Czechia). The F_v_/F_m_ parameter was visualized spatially with Fluorcam 7.0 software according to the protocol provided by Photon Systems Instruments.

## Acknowledgments

We are thankful to Dr. E. Park and Dr. J. Woo for their comments and advice. This work was supported by National Research Foundation of Korea (NRF) grants funded by the Korean government (MSIT) [NRF-2018R1A5A1023599 (SRC), NRF-2021R1A2B5B03001613 and NRF-2019R1C1C1008698].

## Author contributions

J.H.L. and D.C. conceived the experiments. S.L. performed confocal imaging analysis, cell death assays, cRT-PCR, and pathogen virulence assays. J.H.L. performed RIP and iCLIP experiments, protein translation assays, and polysome profiling. J.K. performed plasmid construction, polysome profiling, and statistical analyses. M.K. performed NGS analyses. C.W.M. and S.T.K. assisted LC-MS/MS analyses. S.C. assisted experimental design for cRT-PCR. J.H.L. and D.C. supervised and wrote the manuscript.

## Supplemental figures

**Figure S1.**
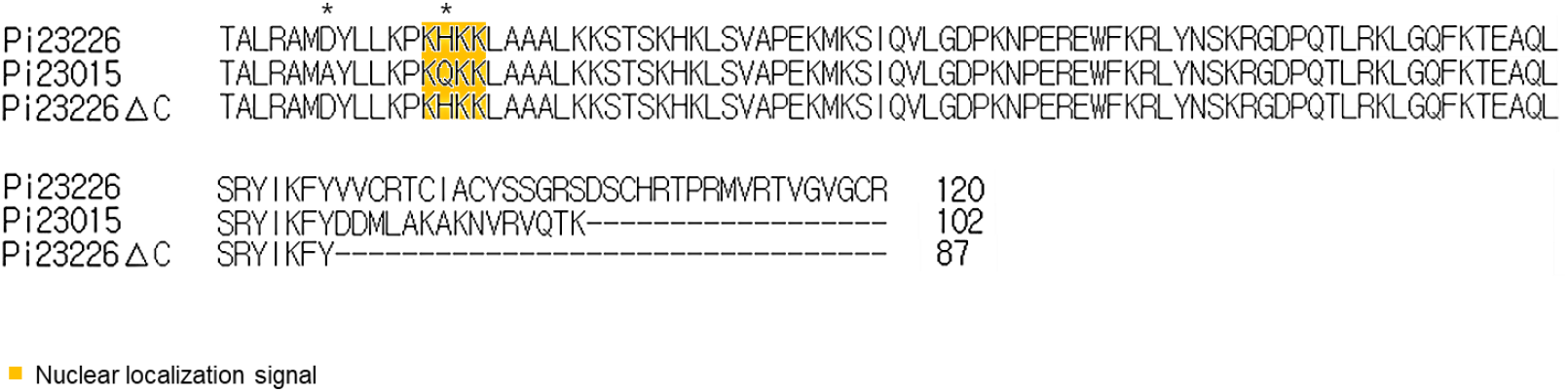
Sequence alignment of Pi23226, Pi23226ΔC, and Pi23015. The sequences of effector domains from each effector was aligned using Color Align Properties (http://www.protocol-online.org/tools/sms2/color_align_prop.html). Residues that are identical or similar biochemical properties were marked with the same color.

**Figure S2.**
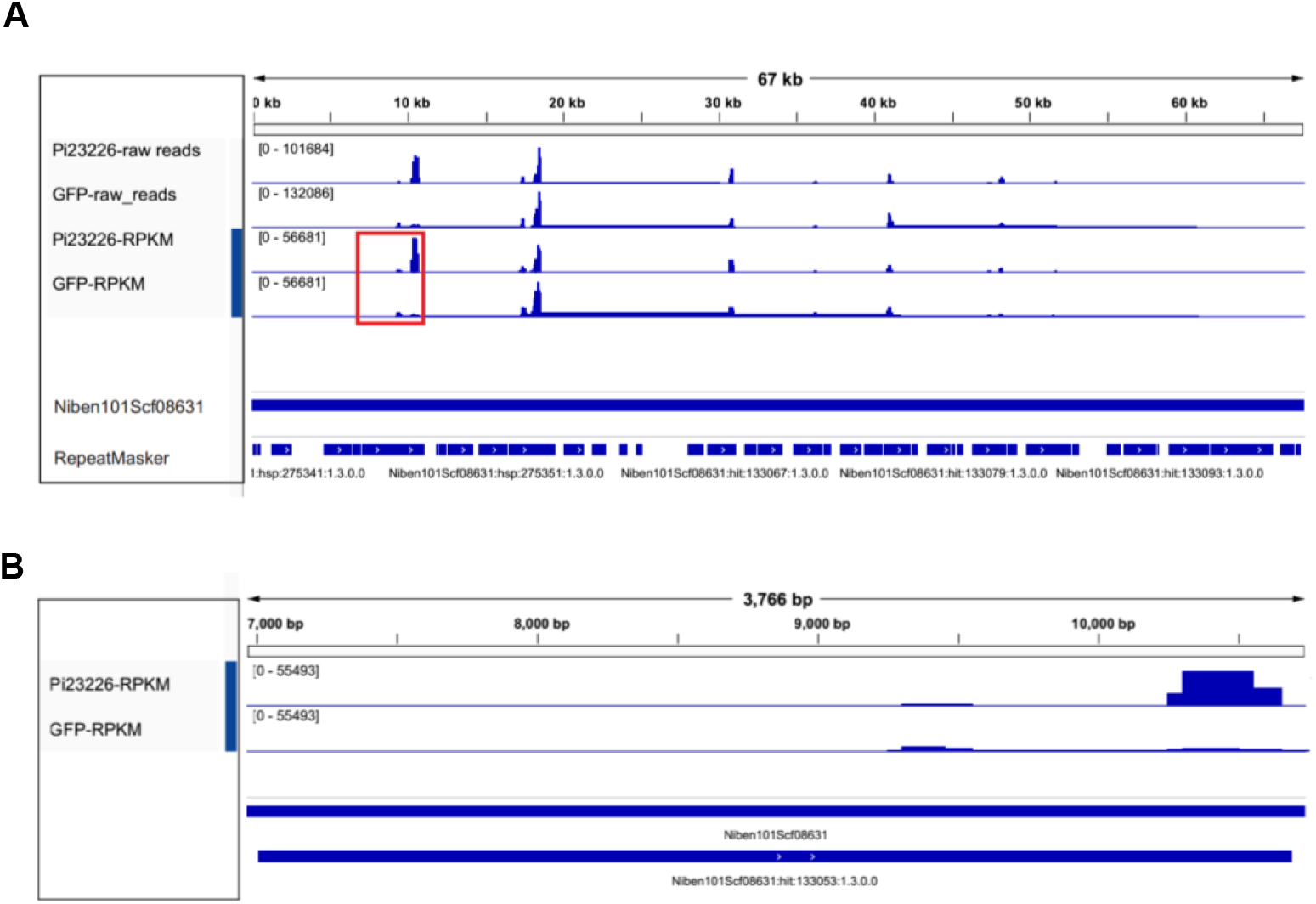
Genomic landscape of the rRNA locus for Pi23226-and GFP-iCLIP analyses (A) The raw and RPKM normalized read coverages were visualized using IGV for the Niben101Scf 08631 scaffold in *N. benthamiana*. (B) Expanded view of red box in (A).

**Figure S3.**
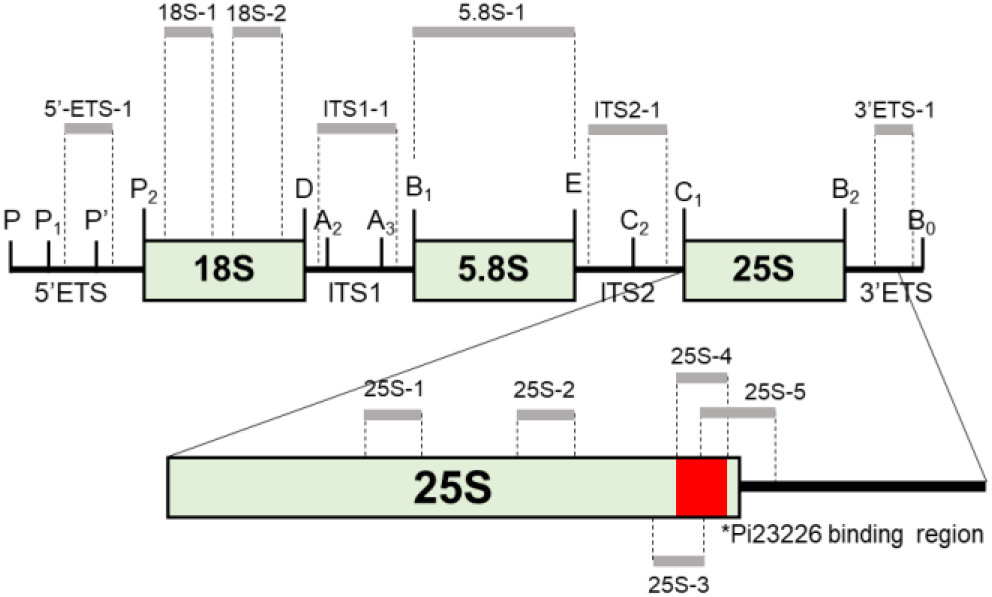
Information of primers used in Figure 2 and 3 was indicated on the 45S rRNA.

**Figure S4.**
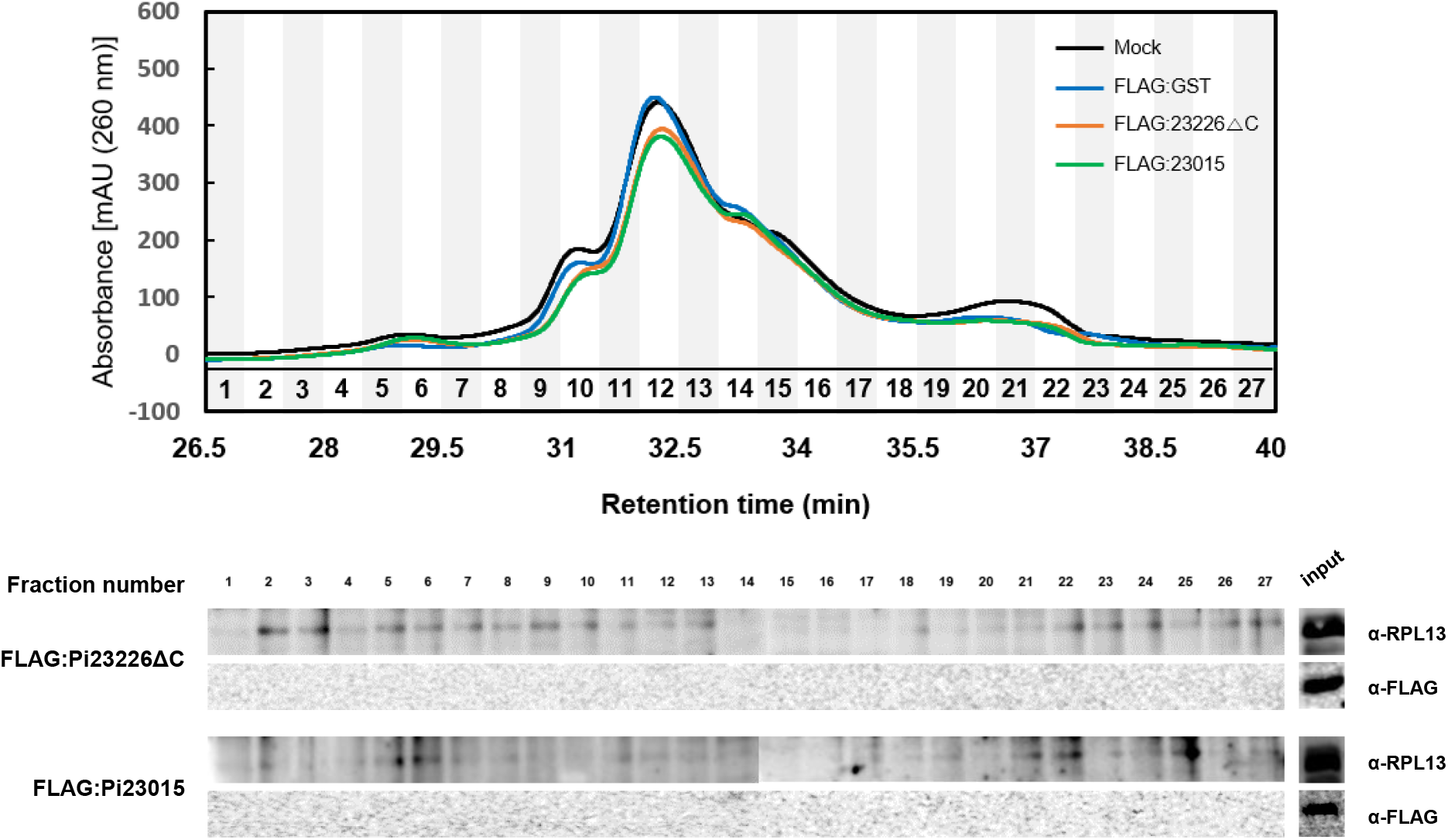
Polysome profiling in FLAG:Pi23226ΔC and Pi23015-expressing *N. benthamiana* using HPLC. Distribution of ribosomes was monitored at absorbance 260nm and fractions were collected in 27 columns with flow rate of 0.4ml/min. Proteins from each fraction was extracted and immunoblots were performed using α-RPL13 or α-FLAG.

**Figure S5.**
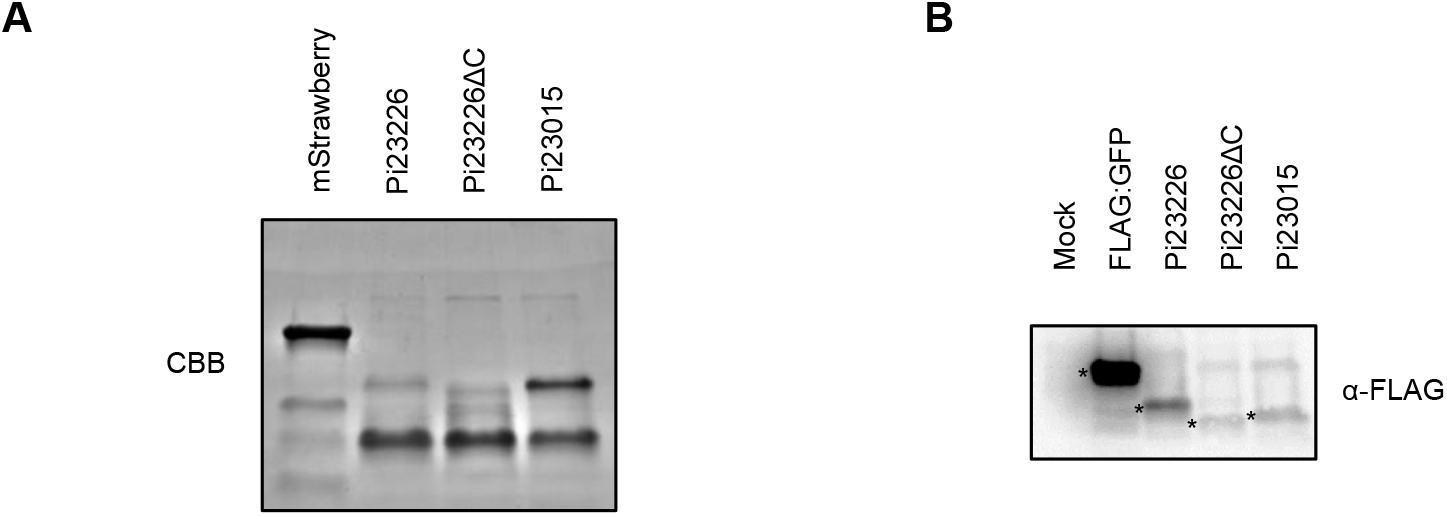
Protein expression of recombinant proteins (A) GST-fused effectors were expressed in *E. coli* (BL21). Recombinant proteins were detected on SDS-PAGE followed by coomassie blue staining. (B) Protein expression of FLAG-effectors in *N. benthamiana*. Leave tissues were collected at 1.5 dpi after agroinfiltration of FLAG-fused effectors. Total protein was extracted and the protein levels of FLAG-effectors were measured using anti-FLAG.

**Figure S6.**
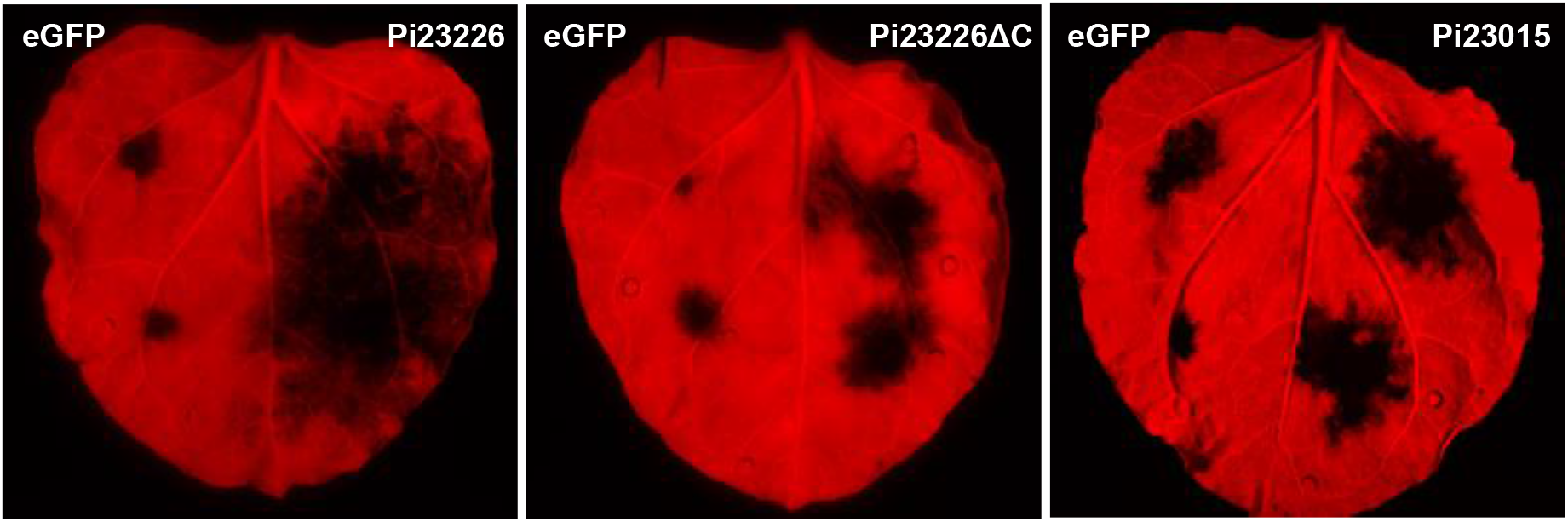
Pi23226 enhances pathogenicity of *P. infestans*. FLAG-effectors were expressed on one-half of the *N. benthamiana* leave and GFP expressed on the other half as a negative control. After 1dpi, zoospores of *P. infestans* (T30-4) was inoculated by dropping (500/drop) on the detached leaves. The lesion size was measured 5 days after inoculation. Each lesion size was photographed using Cy5 and Cy3 channels in Azure 400 (Azure biosystem, USA).

